# Engineered cells that sense and respond to interleukin-23 with neutralizing therapy for autoimmunity

**DOI:** 10.64898/2026.10.12.758578

**Authors:** Gabriela L. Garza, Sonja N. L. Danon, Jonathan Wuchter, Aaron H. Morris

## Abstract

Engineered cell sensors are powerful tools for monitoring biomarkers and delivering timely therapy. Here, we create an engineered cell sensor for interleukin-23 (IL-23), a critical biomarker in many autoimmune diseases, and a therapeutic delivery platform for autoimmune disease management. We utilized Modular Extracellular Sensor Architecture (MESA), a promising platform given its modularity and previous use to developed cell sensors for immune targets such as IL-10. First, we developed a library of IL-23 MESA receptors by incorporating IL-23-specific nanobody binders and performed high-throughput functional screening. We achieved further receptor optimization by modifying the extracellular linker length and the ratio of receptor chain expression. Receptor performance was highly correlated with predicted geometric features such as binder pair distance and degree of transmembrane domain dimerization. The receptors were limited by poor surface expression and functioned most effectively as sensors for intracellular IL-23. Finally, we modified the IL-23 sensing circuit to output an IL-23 neutralizing nanobody with the potential to reduce inflammation triggered by IL-23 secretion. This sense-and-respond architecture can be utilized for dynamic therapeutic treatment to meet the dynamic relapse-remitting pattern of many autoimmune diseases.

**Visal Abstract:** 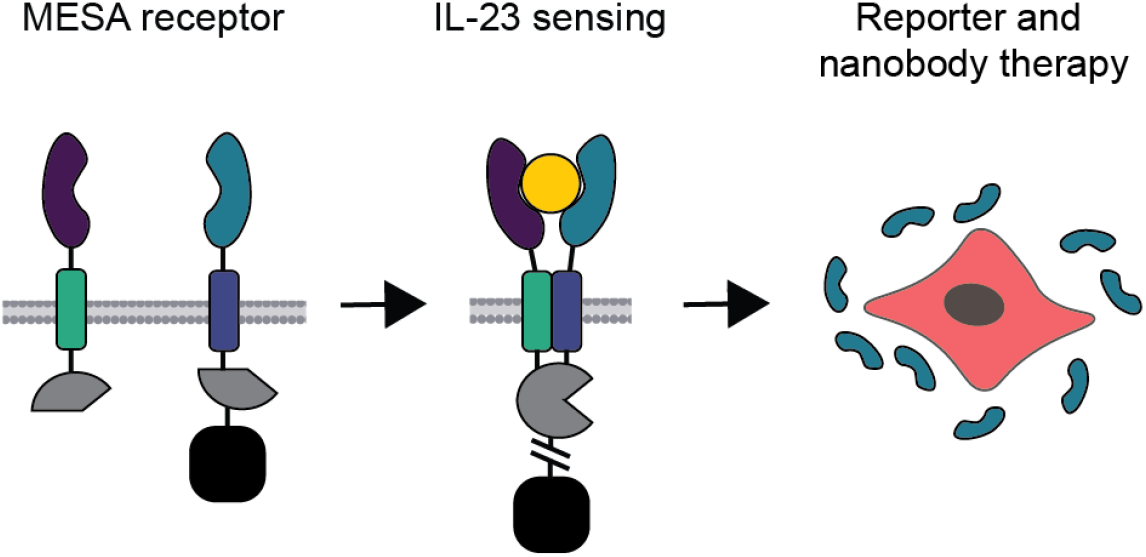

## Introduction

The global incidence of autoimmune disease increases by 12.5% annually, and can be challenging to treat, especially relapsing-remitting forms of disease (e.g. multiple sclerosis (MS), Crohn’s disease, and rheumatoid arthritis (RA)). ^1^ Recent immune modulating therapies have shown promise in treating some of these conditions, but the timing and dose of treatment is patient specific and therefore challenging to control relapses while minimizing total drug delivered and associated side effects. For this reason, effective delivery and timing of treatment remains an unmet challenge in autoimmunity and one that this study aims to address.

The first FDA approved therapies for MS, Crohn’s disease, and RA were approved in 1984, 1993, and 1998 respectively and remained the standard treatments for many years.^2^ Recently, there has been an explosion of newly approved therapies for autoimmune diseases such as upadacitinib, the first oral treatment for Crohn’s disease approved in 2023, risankizumab, an anti-interleukin-23 monoclonal antibody now approved for plaque psoriasis, psoriatic arthritis, Crohn’s disease and ulcerative colitis, and various anti-CD20 antibodies and immunotherapies which have shown great promise in reducing the severity and frequency of relapses in relapsing-remitting MS.^3–6^ These therapies are delivered as daily pills or periodic injections and do not adjust dosage automatically to the changing needs of the patient. If a patient experiences a symptom flare, they may need to contact their doctor to be approved for a higher dose and/or start a round of corticosteroids to get the flare under control.^7^ Delays in this process can cause irreparable tissue damage such as brain lesions in MS patients or joint disfigurement in RA patients.^6,8^ Moreover, these medications can have serious side effects including increased risk of infection.^4,5^ Dynamic approaches to drug delivery may better treat symptoms with the lowest effective dose of drug and thus minimize side effects. An ideal, personalized medicine approach to get the right drug to the right patient at the right time could transform care of autoimmunity, but this remains a challenge with dynamic diseases where different doses of drug may be required at different times.

Living drug factories enable local and timely drug delivery. The earliest example of this is the implantation of encapsulated pancreatic islets which produced insulin and glucagon into a diabetic mouse model published in 1980.^9^ Since then, the field has advanced in terms of biomaterial encapsulation, immunomodulation, and cell circuit engineering, and sent a breakthrough implantable drug delivery device to clinical trials in 2012.^10,11^ A recent report investigated an encapsulated, genetically engineered *E. coli* device that senses a *Pseudomonas aeruginosa* infection and produces an antibiotic to kill it in response.^12^ When co-implanted with a joint prosthetic, the engineered bacteria are able to treat this common type of implant infection locally and adequately despite the poor success rate of treatment with oral or infused antibiotics.^12^ Also, the responsive delivery of antibiotic reduces unnecessary exposure which can cause side effects and contribute to antibiotic resistance.^13^ While bacteria are limited in the complexity of proteins and therefore drugs they can produce, a similar concept can be applied using mammalian cells to create “local drug depots”.^12^ Dr. Omid Veiseh’s group created an encapsulated cytokine-secreting cell device that is co-implanted with human islets to prevent fibrotic capsule formation and protect function of the islets in diabetic animal models.^14^ The continuous delivery of immune-suppressing cytokines eliminated the need for systemic immunosuppressants post-implantation. One limitation of this approach is the lack of a sense-and-respond circuit to regulate drug production. For autoimmune diseases like MS, Crohn’s and RA which often exhibit fluctuating symptom levels, a sense-and-respond circuit that would turn on drug production only when needed would greatly benefit treatment. This way, patients would not receive immunosuppressive therapies when they do not need them.

Interleukin-23 (IL-23) is a key driver of autoimmunity in MS, Crohn’s, RA and others and is an attractive target for sense-and-respond drug delivery. IL-23 is essential for early, local immune responses and plays a key role in organ- or tissue-specific autoimmune diseases.^15^ Specifically, IL-23 is a cytokine that promotes the expansion and maturation of a subset of helper T cells called Th17 cells that produce IL-17 and other pro-inflammatory cytokines.^16^ Ultimately, IL-23 leads to increased inflammation. In fact, mice lacking the gene for IL-23 (p19 subunit) are resistant to developing experimentally induced models of MS and colitis. ^17,18^ In the mouse model of MS, called experimental autoimmune encephalomyelitis (EAE), IL-23 is elevated at the initiation of disease flares and declines as the disease progresses.^19^ In a clinical study, patients with biopsy-confirmed ulcerative colitis had higher levels of serum IL-23 than healthy counterparts, and the level of IL-23 correlated with disease severity.^20^ This supports the rationale that IL-23 could be used as a trigger for therapeutic delivery.

Synthetic receptors are well suited for sensing protein biomarkers. In recent years, the field of oncology has used synthetic receptors to identify and kill cancer cells based on cell surface proteins. This effort was so successful that there are now seven FDA-approved chimeric antigen receptor T (CAR-T) cell therapies for cancer.^21^ Numerous attempts have been made to make CAR-T therapies targeting autoimmune disease including a CAR T-reg therapy targeting cells with overexpressed IL-23 receptors for the treatment of Crohn’s disease, although none have received FDA-approval.^22,23^ In this study, we use a synthetic receptor system that is designed to sense soluble proteins, as opposed to traditional CAR and synthetic Notch receptors which predominantly sense surface-bound ligands. Modular Extracellular Sensor Architecture (MESA) consists of engineered cell surface receptors that trigger a bio-orthogonal cell response upon binding a soluble biological target.^24^ MESA is a highly modular system consisting of two independent receptor chains with a transcription factor (TF) tethered via cleavable linker.^24^ Cleavage of the linker releases the TF and induces a downstream cell function, like the expression of a reporter protein (***Figure 1***).^24^ MESA has been previously developed for various targets including vascular endothelial growth factor (VEGF) and interleukin-10.^25^ In this report, we harness MESA to create an IL-23-responsive therapeutic delivery system for autoimmune disease management.

**Figure 1.**
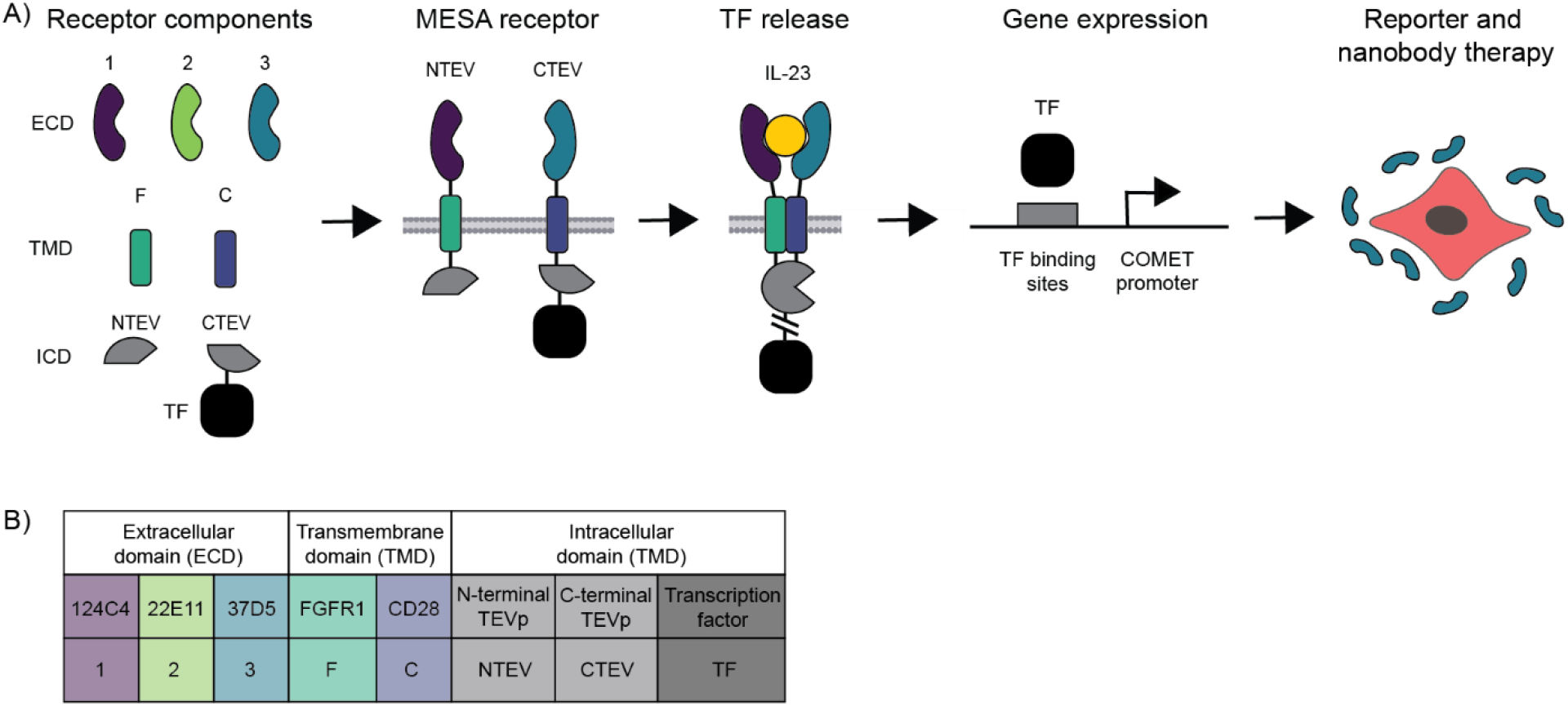
Combinatorial receptor design and responsive drug factory. A) Receptor design using three extracellular domains, two transmembrane domains, and two intracellular domains. IL-23 MESA circuit with an inducible fluorescent reporter and nanobody therapy outputs. B) Legend for receptor components and abbreviations.

## Results

### MESA receptor design

The MESA structure used to detect IL-23 consists of two receptor chains that are each composed of modifiable extracellular domains (ECDs), transmembrane domains (TMDs), and the hallmark split Tobacco Etch Virus protease (TEVp) with attached transcription factor (TF).^27^ When the two chains bind the target ligand, in this case IL-23, they dimerize and the N-terminal and C-terminal fragments of the split protease (NTEV and CTEV respectively) reconstitute to form the active protease.^27^ The reconstituted protease cleaves the polypeptide linker tethering the TF to the CTEV, releasing the TF and enabling it to initiate transcription of its corresponding gene. To achieve bio-orthogonal, tunable cell circuits, MESA receptors were paired with TF and promoter sites from the COMET toolkit.^26,28^ This toolkit has characterized zinc finger transcription factors that avoid genomic binding sites and binding sites of other COMET TF-promoter pairs.^29^ We chose the following three nanobodies as ECDs due to their high binding affinity and strong neutralization of IL-23 and their novelty to the application of engineered cell receptors: 124C4, 22E11, 37D5.^30^ All ECDs were labeled with a Myc protein tag and joined to the TMD with a flexible GS-linker. We used TMDs from fibroblast growth factor receptor 1 (FGFR1) and cluster of differentiation 28 (CD28) as they previously produced high performing MESA receptors.^31^ Using molecular cloning, we created a combinatorial library of 24 receptors in which the two ECDs must bind different epitopes of IL-23 (***Figure 1-2, Table S1***). This paper uses the following receptor nomenclature to indicate the different components used in each respective receptor: TMD and ECD of the NTEV chain, then the TMD and ECD of the CTEV chain (***Figure 1b***). For example, “F2C3” has FGFR1 TMD and nanobody 22E11 in the NTEV chain and CD28 TMD and nanobody 37D5 in the CTEV chain.

**Figure 2.**
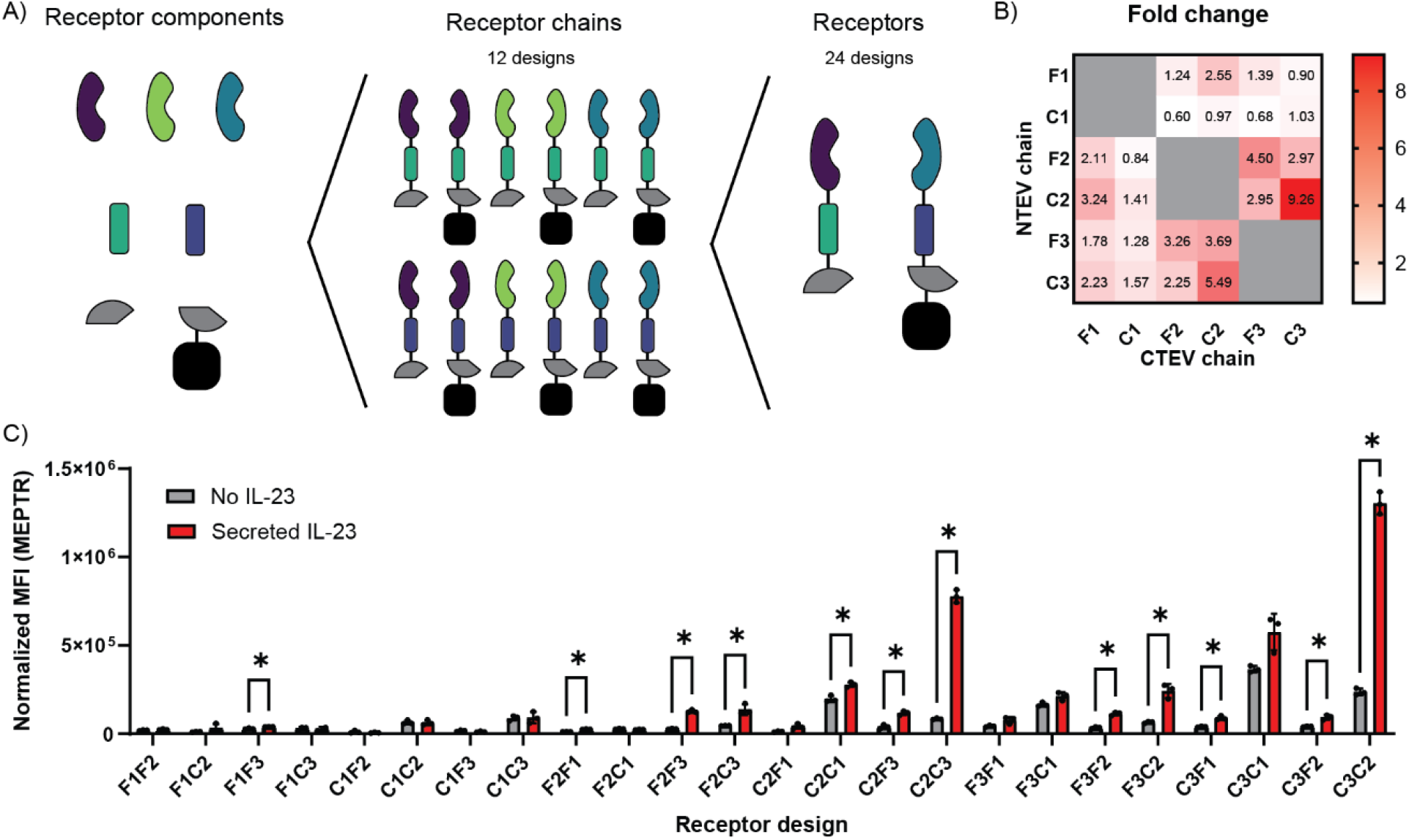
High-throughput functional screening of receptors using co-expressed recombinant human IL-23. A) Graphic illustrating the combinatorial receptor design. B) Heat map of the fold changes of each receptor design. C) Reporter output, DsRedExpress2, of cells transfected with MESA designs with or without co-expressed secreted IL-23. The mean fluorescence intensity of the reporter was converted to molecules of equivalent PE-Texas Red (MEPTR) to normalize between runs, n = 3 biological replicates. *p<0.05

### High-throughput functional assessment of IL-23 MESA receptors

We assessed the ability of each receptor design to detect co-expressed secreted recombinant human IL-23 as an indicator of the receptor’s performance (***Figure 2***). Co-expressed ligand has previously proven to be a robust tool for high-throughput combinatorial screening.^32^ Here we show that 12 of the 24 receptor designs have a significantly higher induced signal compared to background noise. Additionally, receptors with the binder pair #2 and #3 performed better than others. The designs C2C3 and C3C2, both with nanobody binders #2 and #3 and two CD28 TMDs, had the greatest fold changes at 9.26x and 5.49x respectively.

Next, we assessed the ability of our top receptor designs to detect exogenous IL-23 supplemented into the cell media. Unfortunately, none of the receptors produced a detectable induced signal across a range of doses of exogenous recombinant IL-23. To evaluate the potential that cell-secreted ligand exhibited altered folding or glycosylation compared to commercial sources, we transferred conditioned media from cells secreting IL-23 to cells expressing MESA receptors (***Figure S1c***). We determined via ELISA that cells transiently transfected with a secreted IL-23 plasmid produced roughly 1ug/mL IL-23 per day, much higher than we had previously delivered to cells (***Figure S1a***). Of the 16 receptor designs tested, none exhibited a significantly increased signal with conditioned media compared to no ligand.

### Receptor optimization to increase fold change

We employed several strategies to improve the receptor performance measured as fold change. Since IL-23 is a smaller ligand compared to VEGF from which the design was based, we tried decreasing the length of the flexible GS-linker between the ECD and the TMD.^26^ We expected this to improve fold change in the presence of ligand and possibly enable different binding geometries than before. We found that the original length of 32 amino acids was already optimal for receptor C2C3, and that a linker length of 29 amino acids slightly improved the performance of receptor C3C2 (***Figure 3A-B***). We also modified the ratio of receptor chain expression in cells by changing the amount of DNA delivered to cells as has been used to optimize previous MESA receptors (***Figure 3C-D***).^26^ When increasing the expression of NTEV, we noticed a steady decrease in overall signal and fold change. However, the overall signal and fold change were robust when increasing the expression of CTEV from the initial 1:1 ratio.

**Figure 3.**
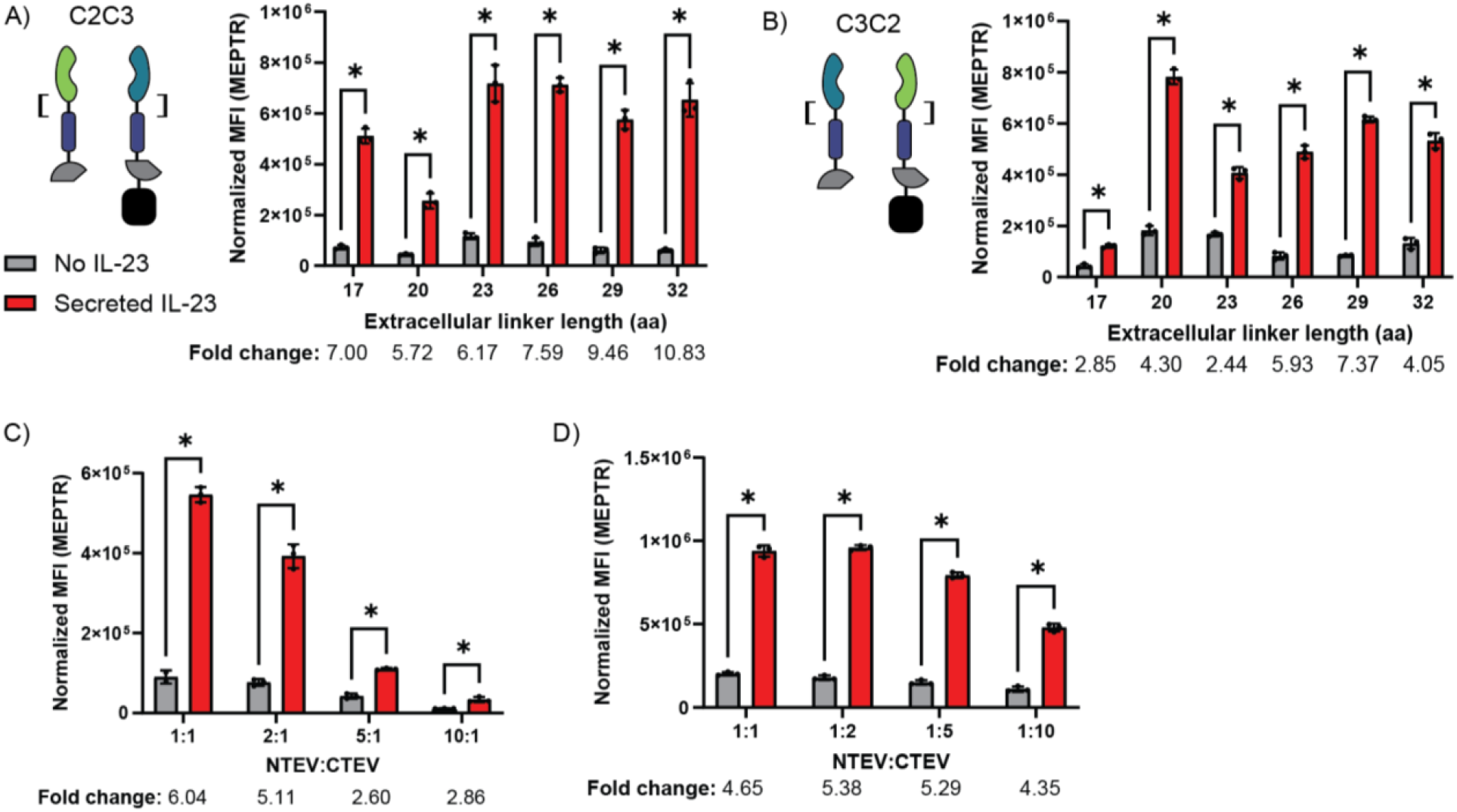
Receptor optimization to increase fold change. A) Reporter output of receptor C2C3 with decreasing extracellular linker length from original 32 aa. B) Reporter output of receptor C3C2 with decreasing extracellular linker length. C) Reporter output of receptor C3C2 with ratiometrically increasing NTEV chain DNA. E) Reporter output of receptor C3C2 with ratiometrically increasing CTEV chain DNA. n = 3 biological replicates for all subplots. *p<0.05

### Predicted geometry analysis

To better understand the design criteria for making a successful synthetic MESA receptor, we performed geometric analyses using the solved crystal structure of IL-23 binding the three nanobodies and predictive protein modeling tools.^30,33^ We generated predicted structures of recombinant IL-23 binding to nanobody pairs using ColabFold (AlphaFold2) and of TMD pairs interacting with each other using PREDDIMER.^34,35^ We used ChimeraX to visualize the structures and take measurements of the ECD distance, TMD distance, TMD contacts, TMD crossing angle, and TMD exit angle of each pair (***Figure 4***).^36,37^ We compared these measurements to experimental values of background reporter signal, secreted IL-23 induced reporter signal, and the fold change between them. The experimental data were log-transformed due to right skew. We found a strong positive correlation between degree of predicted TMD pair interaction (number of contacts and crossing angle) and overall experimental signal (no ligand background and induced signal). We found strong negative correlations between overall experimental signal and other predicted TMD measurements (distance and exit angle). The fold change correlated poorly with all geometric measurements of predicted structures (***Figure 4 & S3***). However, when we perform the same ECD distance measurements using the crystal structure of IL-23 binding the three nanobodies, we find a strong negative correlation between ECD distance and experimental fold change.^30,33^ Essentially, the closer the ECD binders are to each other when binding the ligand, the higher the observed fold change of the receptors with that binder pair.

**Figure 4.**
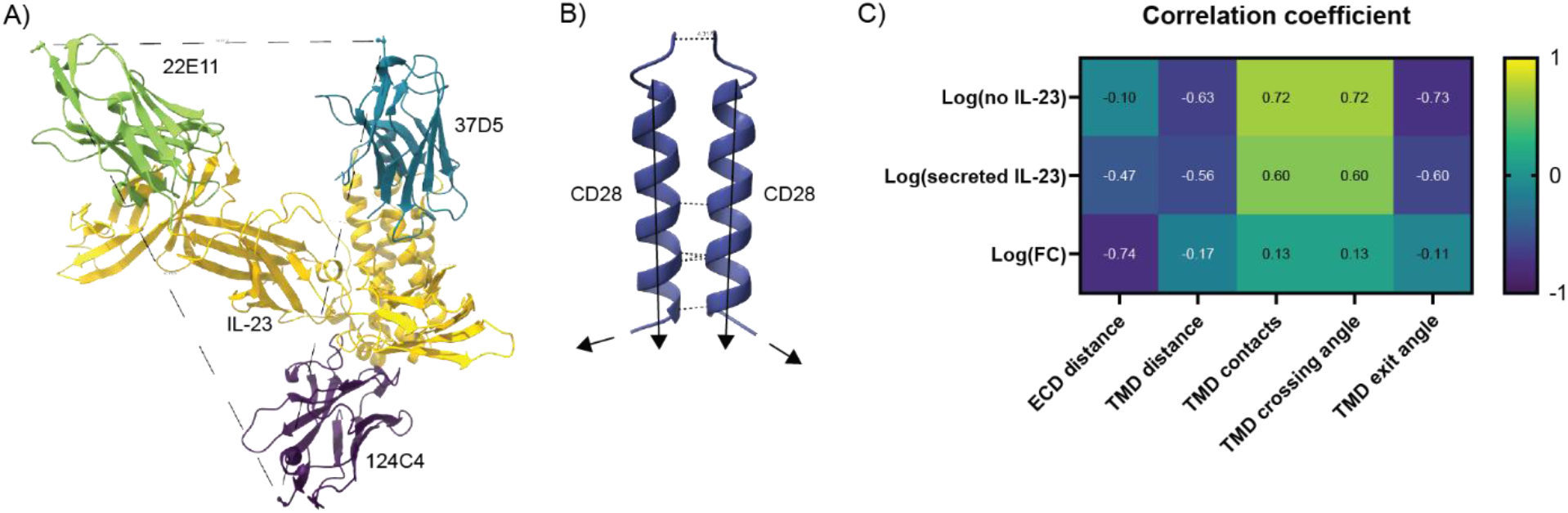
Computational predicted geometry of receptor binding. A) Crystal structure of IL-23 (yellow) binding nanobodies 124C4 (purple), 22E11 (green) and 37D5 (blue). The C-terminal distance is shown here with a dotted black line. B) PREDDIMER prediction of two CD28 TMDs interacting. The TMD contacts and the C-terminus TMD distance are shown with dotted lines. C) Heat map of correlation coefficients between each experimentally observed variable and predicted geometric feature. Experimental values were log-transformed due to right skew.

### Surface expression of receptor chains

We hypothesized that our receptors did not sense exogenous IL-23, despite high inducibility in the presence of co-expressed IL-23, due to poor surface trafficking. We assessed surface trafficking by expressing individual receptor chains in HEK293FT cells and staining with anti-Myc Alexa-647 before analyzing via flow cytometry. Nine of the 12 receptor chains showed significant surface expression compared to a blank plasmid control (***Figure 5***). We performed a multifactorial ANOVA to evaluate the effect of receptor component choice on log-transformed MFI of surface expression. The main effects of ECD and ICD and the following interaction effects significantly impacted receptor surface expression: ECD-TMD, TMD-ICD, and ECD-TMD-ICD. Upon further analysis, receptor chains with 37D5 binders expressed on the surface better than those with 22E11 binders, and CTEV chains expressed better than NTEV chains. To improve surface trafficking, we incorporated the *de novo* protein binder ILP317 into existing designs as an alternate binding domain to our nanobody binders.^38,39^ ILP317 has been found to strongly bind the p19 domain of IL-23 but had not been evaluated as a binder in a synthetic receptor before. We also tested the TMD of the transferrin receptor 1 (TfR) as an alternate component in our receptors to improve surface expression. The TfR has been studied in applications of drug delivery, cancer therapy, and protein engineering.^40^ The version used here is from SpyTag/SpyCatcher protein devices due its strong surface expression and membrane tethering abilities.^41,42^ Unfortunately, altering the ECD and TMD in our receptors resulted in poor surface expression and exhibited low fold-change or even de-inducibility with secreted ligand compared to our original designs (***Figure S2***). Since we found several receptors that sensed co-expressed IL-23, we focused on intracellular sensing applications.

**Figure 5.**
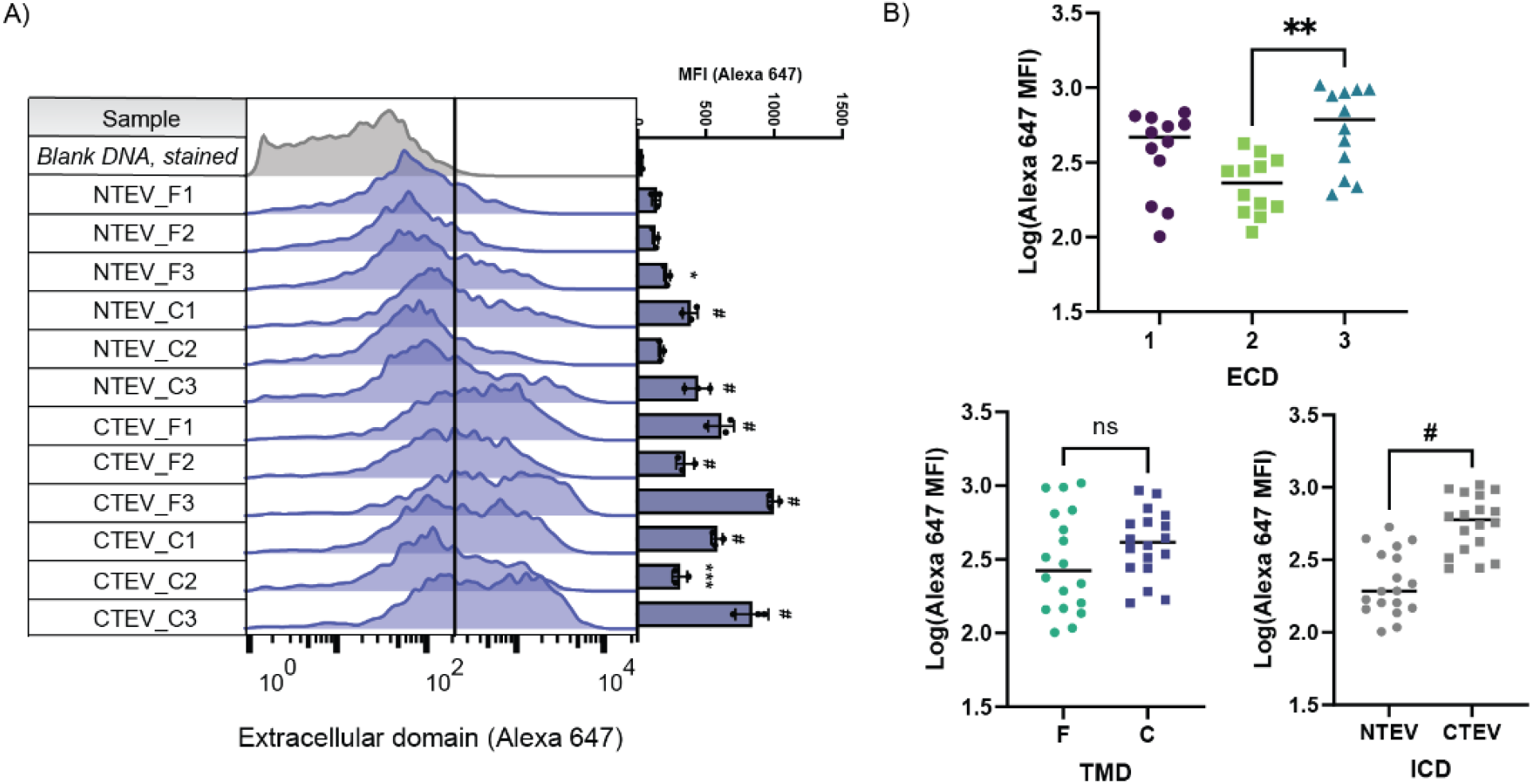
Surface expression of receptor chains. A) Expression of individual receptor chains in HEK293FT cells and stained with anti-Myc fluorescent antibody. Results are shown as fluorescence intensity via flow cytometry. MFI of receptor chain expression for n = 3 biological replicates. Ordinary one-way ANOVA comparing samples to cells transfected with empty vector DNA (blank). B) Subplots of log-transformed MFI of receptor chain surface expression showing main effects of each receptor component, n = 12-18 biological replicates. *p<0.05, **p<0.01, ***p<0.001, ^#^p<0.0001.

### Neutralizing nanobody therapy delivery

To evaluate the therapeutic relevance of our IL-23-sensing cells, we sought to evaluate whether the intracellular sensing machinery enabled us to generate a living drug factory. By coupling the secretion of an IL-23-neutralizing therapy to the cellular expression of IL-23, we reasoned that when a cell occupies a microenvironment in which it will produce IL-23, it would secrete inhibitors to IL-23 in response. This would inactivate IL-23 produced by the engineered cell and mitigate other IL-23 present in the extracellular milieu. The nanobody 37D5 functions effectively as both an IL-23-binding ECD in these MESA receptor designs and has shown to be a strong blocker of IL-23.^30^ To show its efficacy as a cell-delivered therapeutic, we engineered cells to secrete 37D5 upon activation of a MESA receptor by using the same promoter as our fluorescent reporter. This way, the nanobody therapy is released during expression of IL-23, but not in its absence. We co-transfected cells with plasmids encoding receptor C3C2, constitutively secreted IL-23, and zFa6-promoted secretion of 37D5. After 24 hours of incubation, we transferred cell media to a commercially available reporter cell line that produces SEAP in response to IL-23 signaling via native IL-23 receptors (HEK-Blue cells) and measured the IL-23 bioactivity through SEAP production. In ***Figure 6***, we see a dose-dependent (dependent on dose of transfected 37D5 plasmid, pN) decrease in SEAP production of HEK-Blue cells. HEK-Blue cells sense a lower amount of bioactive IL-23 in the presence of the neutralizing nanobody, suggesting that the cell sensors could effectively mitigate IL-23 signaling.

**Figure 6.**
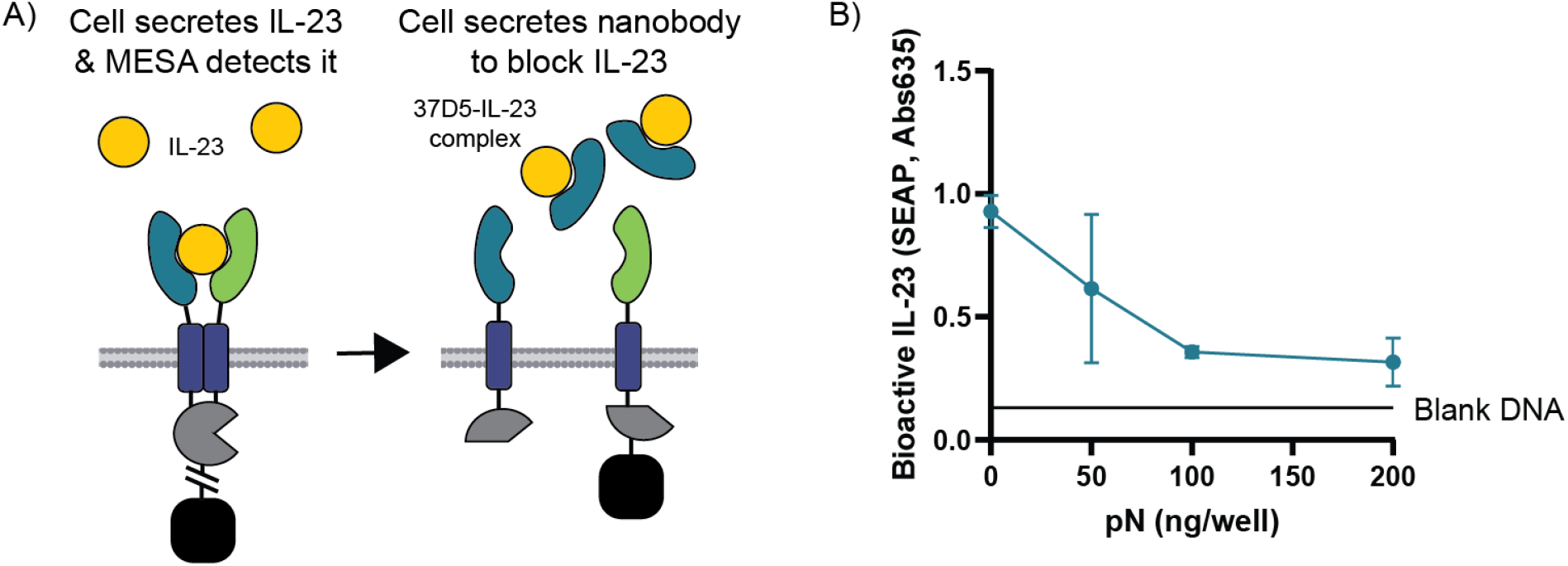
Nanobody therapy delivery neutralizes IL-23. A) Diagram of HEK293FT transfection with MESA receptor, secreted IL23, and varying amounts of secreted 37D5 nanobody plasmid. After expression of transfected cargo, cell media from HEK293FT cells was transferred to HEK-Blue IL23 cells to measure IL23 bioactivity. B) IL-23 bioactivity in the presence of secreted nanobody measured using SEAP secretion from HEK-Blue IL-23 in response to IL-23, n = 3 biological replicates.

## Discussion

Engineered cell sensors have revolutionized medical research by allowing continuous monitoring of disease-specific biomarkers and delivering protein therapies on site and on demand. This is especially impactful to autoimmune disease research and management as many conditions involve fluctuating symptoms. While there have been several attempts to modulate autoimmune inflammation with CAR-T cells, all have focused on surface bound protein targets and fail to take advantage of soluble cell signals like cytokines which are known biomarkers of disease.^23^ In 2015, Schukur et al. created an AND-gated cytokine converter by rewiring natural cytokine receptor outputs to treat psoriasis.^43^ In 2025, Edelstein et al. converted natural cytokine receptors to use bio-orthogonal MESA signaling to create cytokine-dependent gene circuits.^32^ Our group expands the field of engineered cell sensors by using synthetic MESA receptors to sense the cytokine IL-23, the first of its kind and relevant to diseases like MS, Crohn’s, and RA, and deliver a therapeutic output for continuous disease management.

We generated 12 receptors that sense intracellular human IL-23 using specific nanobody binders that had never been used before in a synthetic cell receptor. Previous MESA receptors using nanobody ECDs, GBP2 and GBP7 which bind to GFP, produced a fold change of 9.8x for secreted ligand when incorporated into the split TEVp architecture, which is similar to our findings.^31,44^ Our top performing designs paired 22E11 and 37D5 nanobody binders. A recent study screening AI-generated mini-binders for CARs found that the isoelectric point (pI) of the binder was inversely correlated with surface expression of a receptor chain containing the binder.^45^ We see the same trend for the extracellular binders used in this study (***Figure S3A***) which may help explain why our attempts to improve receptor surface expression by changing the ECD to the *de novo* protein binder ILP317 were unsuccessful. Another consideration is that our receptor constructs are complex and could be experiencing protein misfolding causing aggregation within the cell and therefore lower surface expression.^46^

Our top performers also contained CD28 TMDs on both receptor chains. In previous MESA designs, pairing two CD28 TMDs resulted in high overall signal and low fold change as opposed to high fold change observed in our designs.^31,32^ CARs containing CD28 TMDs have been found to dimerize with natural CD28 receptors inside the cell and reduce surface expression and functionality.^47^ It is reasonable to think that dimerization of MESA receptors containing CD28 TMDs could inhibit surface expression and therefore ability to sense external IL-23. We tried to ameliorate this potential problem by changing the TMD to the stable, well-studied TfR, but were unsuccessful. Focusing on intracellular IL-23 sensing, we further optimized our top receptors by changing the length of the extracellular linkers and the expression ratio of individual receptor chains. These techniques were both used to improve the fold change of a VEGF-sensing MESA receptor.^26^

Prior work investigated predicted structural interactions of natural cytokine receptors containing MESA ICDs which found that shorter ECD and TMD distances, along with greater numbers of ECD and TMD contacts, correlated with higher experimental ligand-induced signal.^37^ When comparing the predicted binding geometry of our receptors to our experimental data, we found that the overall signal (ligand-induced and background) was highly correlated with the number of TMD contacts and crossing angle and inversely correlated with TMD distance and exit angle measured from PREDDIMER predicted binding structures.^35^ We found a strong correlation between high induced signal and fold change versus ECD distance as measured via crystal structure, which is consistent with the results of the VEGF MESA.^26^ We compared this to the ColabFold predicted structures of recombinant IL-23 binding our ECDs with Myc tags and found weaker correlations to our experimental data (***Figure S3B***). This trend held true when we fed the exact same sequences from the crystal structure into ColabFold. Our nanobody binders with Myc tags (135-138 aa) are generally smaller than natural cytokine receptor binders (103-762 aa). However, ColabFold was trained on proteins of average size 268 aa and less than 1,000 aa, so this size range of proteins should be within its predictive limits.^34^ The inclusion of a Myc tag often causes lower confidence in predicted binding for that region, but this does not fully explain why the ColabFold predicted structures differed from the Protein Data Bank structure so much even when using the same sequence.

Finally, we demonstrated how our technology can sense intracellular IL-23 and secrete a neutralizing nanobody therapeutic in response. During the autoimmune inflammatory cascade, IL-23 is primarily secreted by dendritic cells and macrophages.^48,49^ The IL-23-responsive drug factory described here could be implemented in these cell types for early neutralization of IL-23 secreted by the cell and from others within the surrounding microenvironment for autoimmune disease management. An important modification for long-term use would be to integrate drug factory components into the cell genome and link the expression of our MESA construct to the natural secretion of IL-23. An important consideration would be the choice of signal sequence and its impact on MESA receptor trafficking.^50^ In a similar manner to how CAR-T is a sense-and-respond cell therapy, we propose that immune cells could be engineered to respond to inflammatory microenvironments and deliver anti-IL-23 payloads as responsive, living drug factories. The ability to sense-and-respond to autoimmune inflammation at the source could be transformative to the way we treat autoimmunity.

## Methods

### Plasmid generation

All plasmids were generated using PCR and digest-ligate molecular cloning to modify existing plasmids gifted from the lab of Joshua Leonard at Northwestern University. DNA fragments were purchased from Twist, and PCR primers from IDT. All sequences verified through whole-plasmid sequencing through Primordium or Plasmidsaurus.

### Cell culture

The HEK293FT cell line was purchased from ThermoFisher (R70007), and the HEK-Blue IL-23 cell line was purchased from InvivoGen (hkb-il23). The cell line HIE156 containing a genomically integrated DsRedExpress2 fluorescent reporter with zF6 promoter, an engineered form of HEK295FT, was gifted by the Leonard lab.^31^ All cells were tested for mycobacteria upon entering the lab (MP Biomedicals, 0930504-CF). Cells were maintained at a humid 37°C with 5% CO_2_ and passaged at 80-90% confluence. Cells were grown in Dulbecco’s Modified Eagle Medium (DMEM, Sigma-Aldrich, D7777-50L) supplemented with 10% fetal bovine serum (FBS, Avantor, 76419-584), 2 mM L-glutamine (Gibco, 35050061), and 1% Penicillin/Streptomycin (Gibco, 15140-122). HEK-Blue culture medium was also supplemented with 100 ug/mL Normocin for cell maintenance (InvivoGen, ant-nr-50).

### Transfection

Cells were seeded at 1.5x10^5^ cells in 375 uL of complete DMEM per well into 24-well tissue-culture treated plates at the time of transfection. Purified DNA was diluted into OptiMEM (Gibco, 31985-062) and supplemented with pHIE298 filler DNA such that each well received 500 ng of DNA total and 25 uL of DNA-OptiMEM mix. The transfection reagent polyethyleneimine (PEI, Sigma-Aldrich, 408727-100ML) was suspended into sterile water at 1 mg/mL and stored as aliquots at -20°C. PEI aliquots were further diluted into OptiMEM (3.5 uL into 25 uL) at the time of transfection and then mixed 1:1 per volume of DNA-OptiMEM mix. This DNA-PEI-OptiMEM mix was immediately vortexed for 20 seconds and allowed to complex at room temperature for at least 15 minutes. The final transfection mix was pipetted gently, and 50 uL was added dropwise to each well. Plates were rocked gently to mix and incubated overnight. The next morning, cell media was aspirated and replaced with 375 uL/well complete DMEM.

### Interleukin-23 preparation

Recombinant human IL-23 from (Biolegend, 787902) was reconstituted in sterile 1X PBS at 25 mg/mL. IL-23 solution was aliquoted and stored at -80°C. Bioactivity compared to manufacturer-provided sample of hIL-23 confirmed using HEK-Blue IL-23 cells and QUANTI-Blue assay (InvivoGen, rep-qbs).

### Flow cytometry

Working in the dark, transfected cells were harvested 24 hours after media change or ligand addition, approximately 36 hours after transfection. Cells were washed with 1x sterile PBS and either trypsinized (Gibco, 25200-056) for 4-5 minutes at 37°C (functional testing) or incubated at room temperature for 5 minutes with cold 5 mM EDTA FACS buffer (surface expression). Trypsinization was quenched with cold, complete DMEM, and cells were kept on ice from here onward. Cells were washed off plates by pipetting and transferred to a deep 96-well plate. Cells were spun at 200xg and washed with 0.5-1 mL of FACS buffer. For surface expression, cells were stained with anti-Myc Alexa Fluor 647 (BioLegend, 626810) for 20 minutes, then eFluor-780 (Invitrogen, 501129035) for 10 min, then washed three times with FACS buffer. Finally, all cells were resuspended in 250 uL FACS buffer and analyzed via flow cytometry.

We used the BioRad Cell Analyzer ZE5 equipped with 405, 488, 561, and 640 nm lasers. For functional screening, Cells/Single cells/miRFP720+/mNeonGreen+/EBFP2+ gated cells were measured for MFI of DsRedExpress2 for MESA receptors expressed in HIE156 cell line. For surface expression, Cells/Single cells/Live cells/mNeonGreen+ or EBFP2+ gated cells were measured for MFI of Alexa-647. Stop gate of 20,000 single cells and 5% FSC trigger, 1 uL/sec feed rate. See supplementary materials for more cell gating details. To normalize between experiments, we ran UltraRainbow Calibration Particles, 9 Peaks (Spherotech, URCP-100-2H) using the same color voltages as the samples.

### Interleukin-23 bioactivity assay

At least two passages after thawing, HEK-Blue IL-23 cells were seeded in a 96-well tissue culture plate at 5x10^4^ cells/well and filled to 180 uL/well with complete DMEM. Either cell supernatant or recombinant hIL-23 in PBS was added to the seeded cells 20 uL/well and incubated overnight. The next morning, the HEK-Blue plate was sealed, vortexed briefly, and centrifuged at 200xg for 5 minutes. 20 uL of cell supernatant was transferred to a clear, flat-bottom 96-well plate and supplemented with 180 uL/well QUANTI-Blue solution. The plate was immediately placed in a 37°C plate reader (BioTek Cytation 5), and absorbance measured every 30 minutes for 3 hours at 635 nm. See the QUANTI-Blue assay manufacturer protocol from InvivoGen.

### Statistical analysis

Unpaired, two-tailed Student’s t-tests used for signal-noise comparisons. One-way ANOVA and Watson-Tukey conversion used for multiple comparisons. Multifactorial ANOVA with Tukey post-hoc tests were used for comparisons with multiple independent factors.

## Abbreviations

MS: multiple sclerosis
RA: rheumatoid arthritis
IL-23: interleukin-23
CAR: chimeric antigen receptor
MESA: modular extracellular sensor architecture
TF: transcription factor
VEGF: vascular endothelial growth factor
TEVp: tobacco etch virus protease
ICD: intracellular domain
TMD: transmembrane domain
ECD: extracellular domain
FGFR1: fibroblast growth factor receptor 1
CD28: cluster of differentiation 28
NTEV: N-terminal split TEVp
CTEV: C-terminal split TEVp

## Author Information

GLG designed experiments, collected the data, and wrote the manuscript. SNLD provided conceptual and experimental assistance. JW provided experimental assistance. AHM provided funding, designed experiments, performed initial plasmid construction, and edited the manuscript.

## Acknowledgment

This work was supported in part by the NIH Pathway to Independence Award R00EB028840 (A.H.M.), National Institute Of General Medical Sciences Award R35GM162195 (A.H.M.), University of Michigan Rackham Engineering Award (G.G.), and National Institute Of Allergy and Infectious Diseases Award 1F31AI197803-01 (G.G). Any opinions, findings, and conclusions or recommendations expressed in this material are those of the authors and do not necessarily reflect the views of the NIH. We acknowledge the Leonard lab at Northwestern University for gifting us the HIE156 engineered cell line and plasmids for initial library building. We would like to thank the University of Michigan Flow Cytometry core for providing flow cytometry resources.

