## Supplemental information for "Engineered cells that sense and respond to interleukin-23 with neutralizing therapy for autoimmunity"

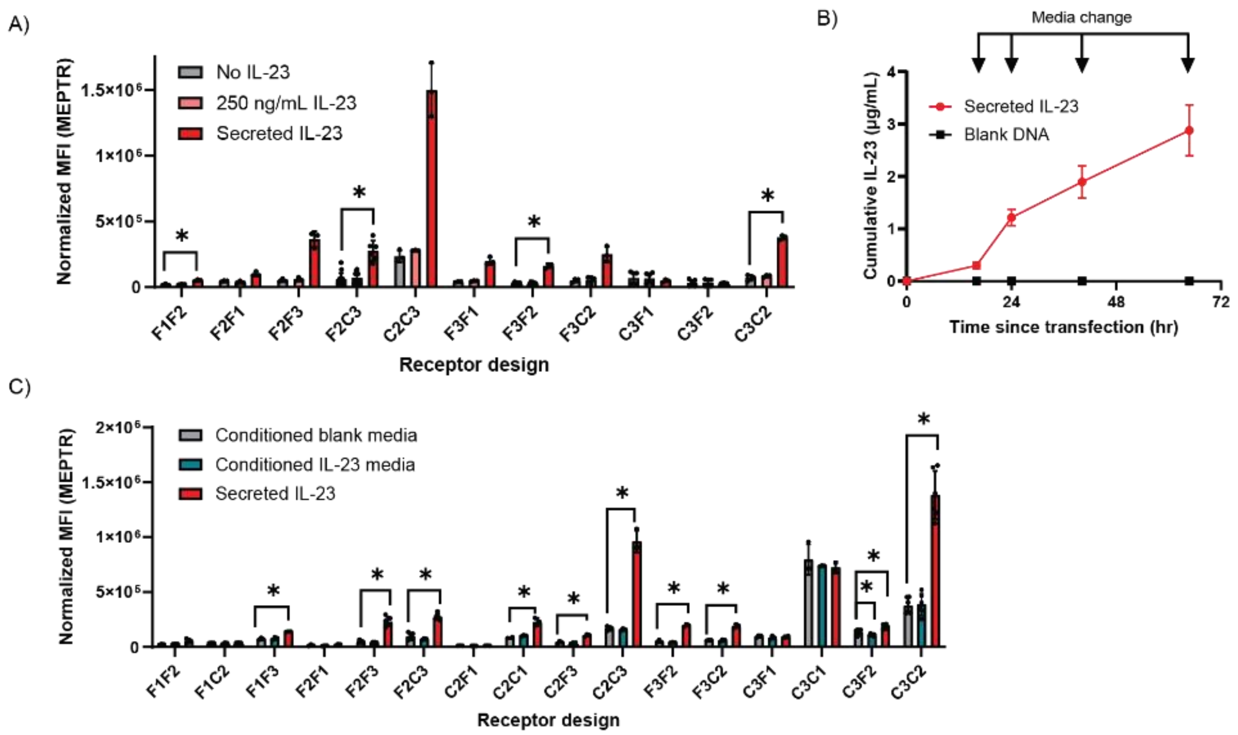


***Figure S1.*** **Exogenous IL-23 supplementation yields no increase in reporter output.** A) Reporter output, DsRedExpress2, of cells transfected with MESA designs with or without co-expressed IL-23, or media supplemented recombinant human IL-23, n = 3-15 biological replicates. B) Cumulative secretion of IL-23 produced by transfected HEK293FT cells measured by ELISA, n = 3 biological replicates. C) Reporter output of cells transfected with MESA designs with conditioned media from cells transfected with blank DNA or secreted IL-23, or co-expressing secreted IL-23, n = 3-6 biological replicates. The mean fluorescence intensity of the reporter was converted to molecules of equivalent PE-Texas Red (MEPTR) to normalize between experiments. *$p<0.05$

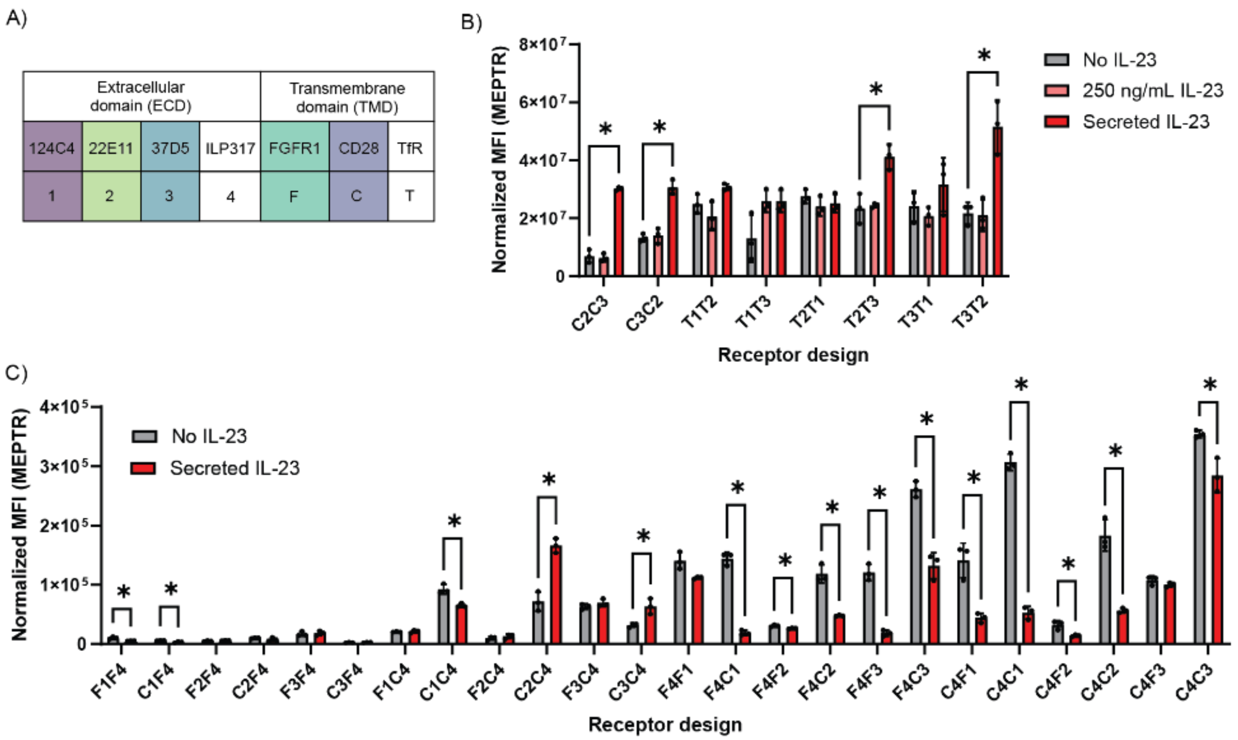


***Figure S2.*** **IL-23 MESA receptors with alternative components.** A) Legend for receptor components. B) Reporter output, DsRedExpress2, of cells transfected with TfR MESA designs with or without co-expressed secreted IL-23, or media supplemented recombinant human IL-23, n = 3 biological replicates. C) Reporter output of cells transfected with ILP317 MESA designs with or without co-expressed IL-23, n = 3 biological replicates. *p<0.05

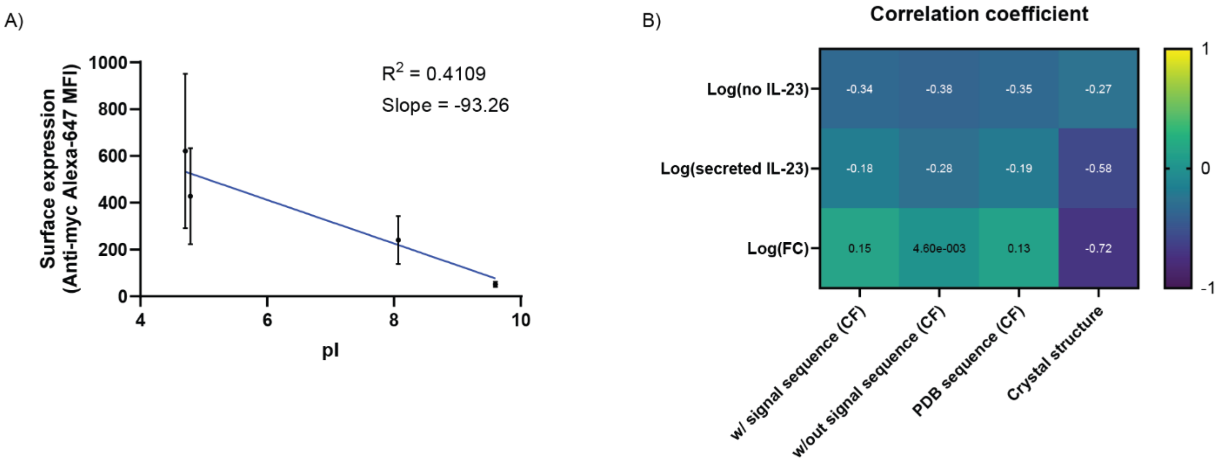


***Figure S3.*** **Structural analysis of MESA receptors.** A) Surface expression of receptor chains measured using anti-Myc Alexa-647 antibody versus isoelectric point of the binder, n = 6-12 biological replicates. B) Heat map of correlation coefficients between each experimentally observed variable and ECD distance. ECD distances were measured from ColabFold (CF) predicted structures using sequences of our ECDs with or without the signal sequence or the published sequence found on the Protein Data Bank. We also compared experimental values to the ECD distances measured from the Protein Data Bank crystal structure. Experimental values were log-transformed due to right skew.

***Table S1. Combinatorial receptor designs for IL-23 MESA.***

| Receptor design | Code name | NTEV chain | | | CTEV chain | | |
|---|---|---|---|---|---|---|---|
| | | **ICD** | **TMD** | **ECD** | **IDC** | **TMD** | **ECD** |
| **1** | F1F2 | NTEV | FGFR1 | 124c4 | CTEV | FGFR1 | 22e11 |
| **2** | F1C2 | NTEV | FGFR1 | 124c4 | CTEV | CD28 | 22e11 |
| **3** | F1F3 | NTEV | FGFR1 | 124c4 | CTEV | FGFR1 | 37D5 |
| **4** | F1C3 | NTEV | FGFR1 | 124c4 | CTEV | CD28 | 37D5 |
| **5** | C1F2 | NTEV | CD28 | 124c4 | CTEV | FGFR1 | 22e11 |
| **6** | C1C2 | NTEV | CD28 | 124c4 | CTEV | CD28 | 22e11 |
| **7** | C1F3 | NTEV | CD28 | 124c4 | CTEV | FGFR1 | 37D5 |
| **8** | C1C3 | NTEV | CD28 | 124c4 | CTEV | CD28 | 37D5 |
| **9** | F2F1 | NTEV | FGFR1 | 22e11 | CTEV | FGFR1 | 124c4 |
| **10** | F2C1 | NTEV | FGFR1 | 22e11 | CTEV | CD28 | 124c4 |
| **11** | F2F3 | NTEV | FGFR1 | 22e11 | CTEV | FGFR1 | 37D5 |
| **12** | F2C3 | NTEV | FGFR1 | 22e11 | CTEV | CD28 | 37D5 |
| **13** | C2F1 | NTEV | CD28 | 22e11 | CTEV | FGFR1 | 124c4 |
| **14** | C2C1 | NTEV | CD28 | 22e11 | CTEV | CD28 | 124c4 |
| **15** | C2F3 | NTEV | CD28 | 22e11 | CTEV | FGFR1 | 37D5 |
| **16** | C2C3 | NTEV | CD28 | 22e11 | CTEV | CD28 | 37D5 |
| **17** | F3F1 | NTEV | FGFR1 | 37D5 | CTEV | FGFR1 | 124c4 |
| **18** | F3C1 | NTEV | FGFR1 | 37D5 | CTEV | CD28 | 124c4 |
| **19** | F3F2 | NTEV | FGFR1 | 37D5 | CTEV | FGFR1 | 22e11 |
| **20** | F3C2 | NTEV | FGFR1 | 37D5 | CTEV | CD28 | 22e11 |
| **21** | C3F1 | NTEV | CD28 | 37D5 | CTEV | FGFR1 | 124c4 |
| **22** | C3C1 | NTEV | CD28 | 37D5 | CTEV | CD28 | 124c4 |
| **23** | C3F2 | NTEV | CD28 | 37D5 | CTEV | FGFR1 | 22e11 |
| **24** | | NTEV | CD28 | 37D5 | CTEV | CD28 | 22e11 |
| **25** | F1F4 | NTEV | FGFR1 | 124c4 | CTEV | FGFR1 | ILP317 |
| **26** | C1F4 | NTEV | CD28 | 124c4 | CTEV | FGFR1 | ILP317 |
| **27** | F2F4 | NTEV | FGFR1 | 22e11 | CTEV | FGFR1 | ILP317 |
| **28** | C2F4 | NTEV | CD28 | 22e11 | CTEV | FGFR1 | ILP317 |
| **29** | F3F4 | NTEV | FGFR1 | 37D5 | CTEV | FGFR1 | ILP317 |
| **30** | C3F4 | NTEV | CD28 | 37D5 | CTEV | FGFR1 | ILP317 |
| **31** | F1C4 | NTEV | FGFR1 | 124c4 | CTEV | CD28 | ILP317 |
| **32** | C1C4 | NTEV | CD28 | 124c4 | CTEV | CD28 | ILP317 |
| **33** | F2C4 | NTEV | FGFR1 | 22e11 | CTEV | CD28 | ILP317 |
| **34** | C2C4 | NTEV | CD28 | 22e11 | CTEV | CD28 | ILP317 |
| **35** | F3C4 | NTEV | FGFR1 | 37D5 | CTEV | CD28 | ILP317 |
| **36** | C3C4 | NTEV | CD28 | 37D5 | CTEV | CD28 | ILP317 |
| **37** | F4F1 | NTEV | FGFR1 | ILP317 | CTEV | FGFR1 | 124c4 |
| **38** | F4C1 | NTEV | FGFR1 | ILP317 | CTEV | CD28 | 124c4 |
| **39** | F4F2 | NTEV | FGFR1 | ILP317 | CTEV | FGFR1 | 22e11 |
| **40** | F4C2 | NTEV | FGFR1 | ILP317 | CTEV | CD28 | 22e11 |
| **41** | F4F3 | NTEV | FGFR1 | ILP317 | CTEV | FGFR1 | 37D5 |
| **42** | F4C3 | NTEV | FGFR1 | ILP317 | CTEV | CD28 | 37D5 |
| **43** | C4F1 | NTEV | CD28 | ILP317 | CTEV | FGFR1 | 124c4 |
| **44** | C4C1 | NTEV | CD28 | ILP317 | CTEV | CD28 | 124c4 |
| **45** | C4F2 | NTEV | CD28 | ILP317 | CTEV | FGFR1 | 22e11 |

| | | | | | | | |
|---|---|---|---|---|---|---|---|
| **46** | C4C2 | NTEV | CD28 | ILP317 | CTEV | CD28 | 22e11 |
| **47** | C4F3 | NTEV | CD28 | ILP317 | CTEV | FGFR1 | 37D5 |
| **48** | C4C3 | NTEV | CD28 | ILP317 | CTEV | CD28 | 37D5 |
| **49** | T1T2 | NTEV | TfR | 124c4 | CTEV | TfR | 22e11 |
| **50** | T1T3 | NTEV | TfR | 124c4 | CTEV | TfR | 37D5 |
| **51** | T1T4 | NTEV | TfR | 124c4 | CTEV | TfR | ILP317 |
| **52** | T2T1 | NTEV | TfR | 22e11 | CTEV | TfR | 124c4 |
| **53** | T2T3 | NTEV | TfR | 22e11 | CTEV | TfR | 37D5 |
| **54** | T2T4 | NTEV | TfR | 22e11 | CTEV | TfR | ILP317 |
| **55** | T3T1 | NTEV | TfR | 37D5 | CTEV | TfR | 124c4 |
| **56** | T3T2 | NTEV | TfR | 37D5 | CTEV | TfR | 22e11 |
| **57** | T3T4 | NTEV | TfR | 37D5 | CTEV | TfR | ILP317 |
| **58** | T4T1 | NTEV | TfR | ILP317 | CTEV | TfR | 124c4 |
| **59** | T4T2 | NTEV | TfR | ILP317 | CTEV | TfR | 22e11 |
| **60** | T4T3 | NTEV | TfR | ILP317 | CTEV | TfR | 37D5 |
